# UBE2N is essential for maintenance of skin homeostasis and suppression of inflammation

**DOI:** 10.1101/2023.12.01.569631

**Authors:** Min Jin Lee, Manel Ben Hammouda, Wanying Miao, Arinze Okafor, Yingai Jin, Huiying Sun, Vaibhav Jain, Vadim Markovtsov, Yarui Diao, Simon G. Gregory, Jennifer Y. Zhang

## Abstract

UBE2N, a Lys63-ubiquitin conjugating enzyme, plays critical roles in embryogenesis and immune system development and function. However, its roles in adult epithelial tissue homeostasis and pathogenesis are unclear. We generated conditional mouse models that deleted *Ube2n* in skin cells in a temporally and spatially controlled manner. We found that *Ube2n-* knockout (KO) in the adult skin keratinocytes induced a range of inflammatory skin defects characteristic of psoriatic and actinic keratosis. These included eczematous inflammation, epidermal and dermal thickening, parakeratosis, and increased immune cell infiltration, as well as signs of edema and blistering. Single cell transcriptomic analyses and RT-qPCR showed that *Ube2n* KO keratinocytes expressed elevated myeloid cell chemo-attractants such as *Cxcl1* and *Cxcl2* and decreased the homeostatic T lymphocyte chemo-attractant, *Ccl27a*. Consistently, the infiltrating immune cells of *Ube2n*-KO skin were predominantly myeloid-derived cells including neutrophils and M1-like macrophages that were highly inflammatory, as indicated by expression of *Il1β* and *Il24.* Pharmacological blockade of the IL-1 receptor associated kinases (IRAK1/4) alleviated eczema, epidermal and dermal thickening, and immune infiltration of the *Ube2n* mutant skin. Together, these findings highlight a key role of keratinocyte-UBE2N in maintenance of epidermal homeostasis and skin immunity and identify IRAK1/4 as potential therapeutic target for inflammatory skin disorders.

## INTRODUCTION

Ubiquitination is a reversible post-translational modification that regulates protein stability and function (1). In general, ubiquitination is a stepwise process that include activation of a ubiquitin (Ub) monomer by an E1 enzyme (such as UBA1) and transfer of the activated Ub to an E2 conjugating enzyme, and subsequently onto the ε-amino group of a lysine residue of the substrate or the preceding Ub (1–3). The N-terminal methionine residue (M1) and each of the seven lysine residues (K6, K11, K27, K29, K33, K48, and K63) of the 76-amino acids Ub polypeptide can be used for attachment of a subsequent Ub, forming structurally and functionally distinct mono-, multi-, or poly-ubiquitin chains (1–3). Ubiquitination is subject to negative regulation by deubiquitinases (DUBs) such as CYLD which removes K63-Ub and M1-Ub from target proteins, and thereby negatively regulates NF-κB and JNK pathways (4, 5).

UBE2N (also known as UBC13) is a K63-Ub-specific E2 conjugating enzyme. Unlike most other E2s, UBE2N is not known for adding Ub directly onto target proteins (6–8). Instead, UBE2N heterodimerizes with a catalytically inactive variant partner, UBE2V1 or UBE2V2, and extends subsequent linkage onto substrates that already contain mono-Ub or other linkages (6–8). UBE2N/UBE2V1 (also known as UEV1A) heterodimers together with TRAF6 as the E3 ligase mediate K63-Ub polyubiquitination and activation of the NF-κB and MAPK signaling pathway (9–14). UBE2N/UBE2V2 (also known as MMS2) heterodimers participate in K63-Ub primarily in nucleus and regulate DNA repair and cell cycle (8, 15–17).

Balanced activity of ubiquitinates and DUBs is crucial for human health. Somatic loss-of-function mutation of UBA1, the major E1 enzyme located on the X chromosome, results in treatment-refractory and often fatal inflammatory syndrome in adult males that includes neutrophilic cutaneous inflammation (18). Similarly, UBE2N loss-of-function is associated with many immunological and neuro-degenerative disorders, as well as embryonic developmental defects of the epidermis (9, 19–22). Conversely, UBE2N overexpression is detected in cancer cells, and it promotes the growth of various cancers, such as neuroblastoma, acute myeloid leukemia, breast cancer, colon cancer, and melanoma (23–28). In contrast, CYLD is well-known as a tumor suppressor whose loss-of-function is implicated in melanoma and nonmelanoma skin cancers, as well as psoriasis (29–32), further highlighting the importance of K63-Ub-mediated signaling in skin health.

Skin serves as a physical and immune barrier providing the first line of defense against many external insults such as UV damage, chemicals, allergens, and pathogenic microbes. Maintenance of skin homeostasis and proper regeneration upon damage are critical for a sustained defense against external insults, as well as inside-out barrier functions. Dysregulation of skin homeostasis is linked to many cutaneous and systemic inflammatory diseases such as atopic dermatitis, psoriasis, and systemic lupus erythematosus (SLE) (33–35). To that end, a previous study has reported that *Krt5*-Cre-mediated keratinocyte-specific deletion of *Ube2n* leads to abnormal development of the skin accompanied by postnatal lethality (22), prohibiting analysis of UBE2N function in adult skin. The *Ube2n* knockout newborn epidermis shows defects in differentiation, decreased proliferation, and increased apoptosis (22). While these data underscore the importance of UBE2N in embryonic epidermal development and postnatal homeostasis, the role of UBE2N in adult skin homeostasis is yet to be determined.

To address this gap in the field, we utilized two conditional *Ube2n* knockout (KO) mouse models: the ubiquitous *Rosa*26^CreERT2^ and the basal epidermal keratinocyte-specific *Krt5*^CreERT2^ transgenic lines that permit inducible gene targeting in adult skin via topical treatments of 4-hydroxy-tamoxifen. We found that topically induced deletion of *Ube2n* in adult skin leads to inflammatory skin lesions marked by eczema, epidermal and dermal thickening, parakeratosis, and increased immune cell infiltration. We also found that loss of UBE2N in keratinocytes is sufficient to induce inflammatory skin defects. The *Ube2n* KO keratinocytes produced elevated levels of pro-inflammatory cytokine genes such as *Il1β* and *Il24* as well as myeloid cell chemokines such as *Cxcl1* and *Cxcl2*. In contrast, *Ccl27a*, that encodes a regulator of immune homeostasis, is downregulated (36). Consistently, the mutant skin contained abundant infiltration of myeloid cells, including neutrophils and M1-skewed macrophages. Furthermore, we found that the interleukin-1 receptor-associated kinase 1 and 4 (IRAK1/4) was highly activated in the mutant skin, and treatment of an IRAK1/4-specific pharmacological inhibitor dampened the inflammatory phenotypes. Our findings reveal an indispensable role of UBE2N in maintaining epidermal homeostasis and suppression of skin inflammation and identify IRAK1/4 signaling pathway as a potential therapeutic target for inflammatory skin disorders.

## RESULTS

### Deletion of UBE2N in skin leads to inflammation

To investigate the role of UBE2N in adult skin, we first utilized *Rosa26*^CreER^.*Ube2n*^fl/fl^ mice, in which ubiquitous expression of Cre recombinase allows temporally and spatially localized deletion of *Ube2n* via topical treatments of 4-hydroxytamoxifen (4-OHT) (Figure 1A). We observed that deletion of *Ube2n* in adult back skin via topical treatments of 4-OHT led to the development of a range of skin abnormalities, including eczema with crusted erosions that resembled psoriatic plaques, as well as skin blistering in some cases (Figure 1B). Histological analysis of the mutant skin collected 2-3 weeks post-4-OHT treatments revealed abnormal hair growth, hyperkeratosis, thickening of the epidermal and dermal compartments, and markedly increased immune cell infiltration (Figure 1C). Interestingly, one-time brief exposure of low dose 4-OHT induced initial inflammation on *Rosa26^CreER^*.*Ube2n*^fl/fl^ back skin around 2 weeks after exposure (Figures S1A-B). However, this inflammation was resolved over time (2-3 weeks later) with normal hair growth (Figure S1C). This suggests that extensive deletion of *Ube2n* in the skin is required to induce progressive and irreversible epidermal abnormalities and inflammation.

**Figure 1.**
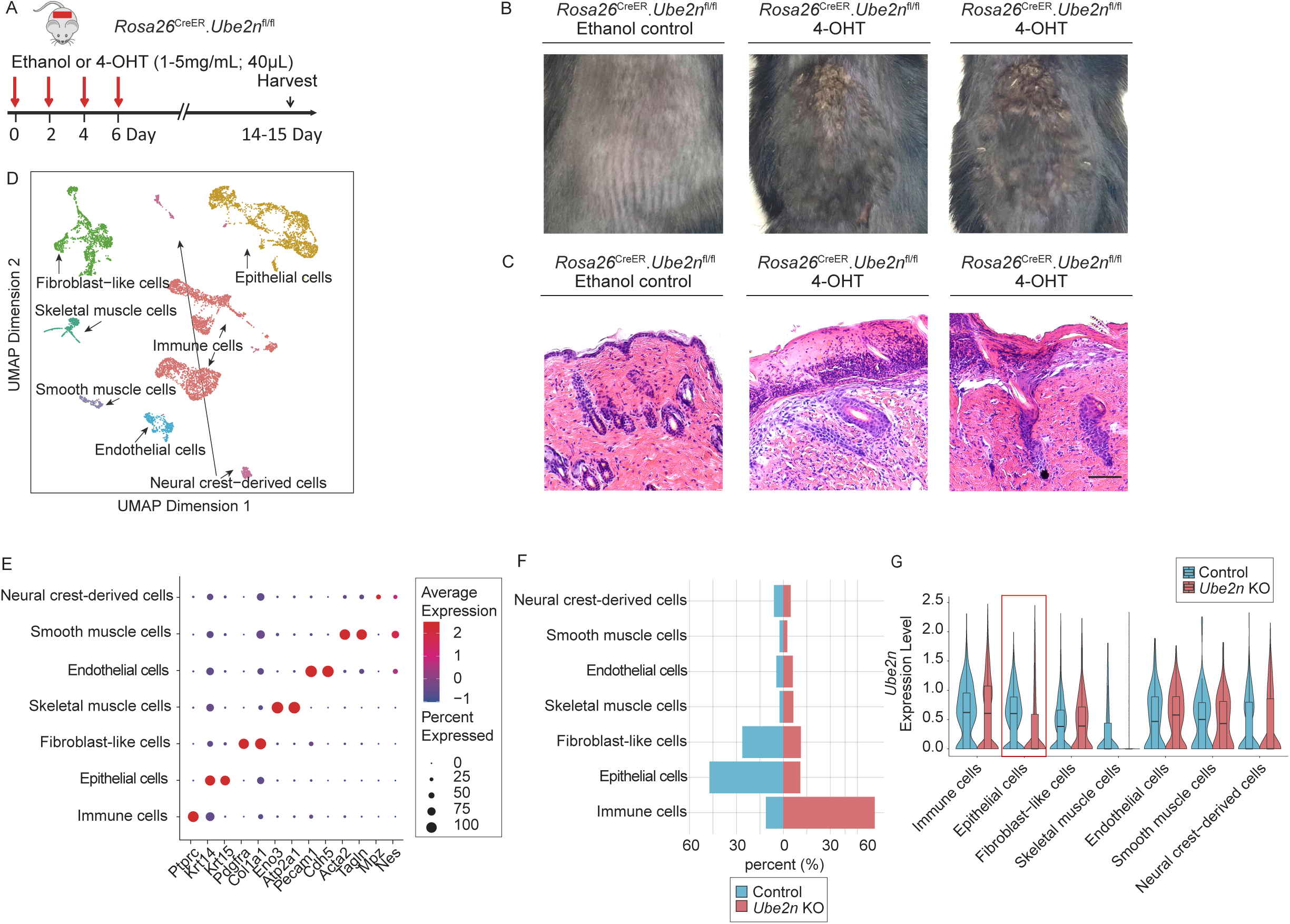
Topically induced deletion of *Ube2n* in the skin leads to inflammation. A. Experimental scheme of induction of *Ube2n* deletion in 6-10 weeks old *Rosa26*^CreER^.*Ube2n*^fl/fl^ mice. 40μL of 5 mg/mL of 4-hydroxytamoxifen (4-OHT) was applied topically every 48 hours for four times to ensure deletion of *Ube2n*. B. Appearance of mouse back skin 15 days post-induction. C. H&E histological analysis of the mouse skin sections. Scale bar: 100μm. D. UMAP of skin cells identifying 7 cell populations. Single-cell RNA-seq analysis of control and *Ube2n* KO skin samples (n=2-3 / group pooled). E. Dot plot of cell identification with key genes. F. Percent distribution of the control and the *Ube2n* KO skin cells. G. Violin and box plot of *Ube2n*.

### Single-cell RNA-seq reveals immune infiltration in the UBE2N mutant skin and various gene expression changes in UBE2N deficient keratinocytes

To understand mechanisms of UBE2N of regulation of skin homeostasis, we performed single-cell RNA-sequencing (scRNA-seq) analysis of the control skin samples and the mutant skin samples of adult 9 weeks-old *Rosa26*^CreER^.*Ube2n*^fl/fl^ mice collected 2 weeks after treatment of vehicle control or 4-OHT. A total of 4147 of control and 4586 of mutant skin cells were sequenced and passed quality control. Through unbiased clustering of the datasets, we identified 7 broad cell types (Figure 1D): epithelial cells (*Krt14*^+^, *Krt15*^+^); immune cells [*Ptprc*^+^ (CD45)]; fibroblast-like cells (*Col1a1*^+^, *Pdgfra*^+^); skeletal muscle cells (*Eno3*^+^, *Atp2a1*^+^); endothelial cells (*Pecam1^+^*, *Cdh5^+^*); smooth muscle cells (*Acta2^+^*, *Tagln^+^*); and neural crest-derived cells (*Mpz*^+^, *Nes*^+^) (Figure 1E). Percent distribution of the cell population revealed that there was a stark increase in the number of immune cells in the *Ube2n* KO skin (Figure 1F), a trend that agreed with our histological analysis (Figure 1C). Significantly decreased expression of *Ube2n* was detected only in epithelial cells of the KO skin compared to control counterparts and no significant changes were observed in other cell types, including fibroblasts, immune cells, and endothelial cells (Figure 1G). Skeletal muscle cells and neural crest-derived cells generally expressed low levels of *Ube2n* indicated by the absence of a visible median line in the boxplot (Figure 1G). These data indicated that the topical application of 4-OHT was effective in targeting epidermal cells, but ineffective in reaching cells in the dermis.

To understand the gene expression changes in epidermal keratinocytes upon *Ube2n* deletion, we further analyzed keratinocytes at a higher resolution and identified 11 subclusters (Figures 2A-C). As expected, *Ube2n* expression was reduced in the KO keratinocytes (Figure 2D). Interfollicular keratinocytes (KC) including basal KC (*Krt14*^+^, *Mt2*^+^), suprabasal KC (termed differentiated KC; *Krt10*^+^, *Krt1*^+^), and terminally differentiated KC (termed keratinized KC; *Lor*^+^, *Flg2*^+^, *Ivl*^+^) showed particularly low expression of *Ube2n* (Figures 2B-D). Hair follicle (HF) cell populations such as upper HF II and bulge I-IV cells did not show lower expression of *Ube2n* (Figure 2D), further supporting the notion that the topical application of 4-OHT effectively deletes *Ube2n* in the cells located in the epidermis, but not those in the dermis. In addition, we identified two clusters of inflammatory KC groups (inflammatory KC I and inflammatory KC II) that expressed high levels of *Krt6a*, *Krt6b*, and *Krt16* (Figures 2A-C).

**Figure 2.**
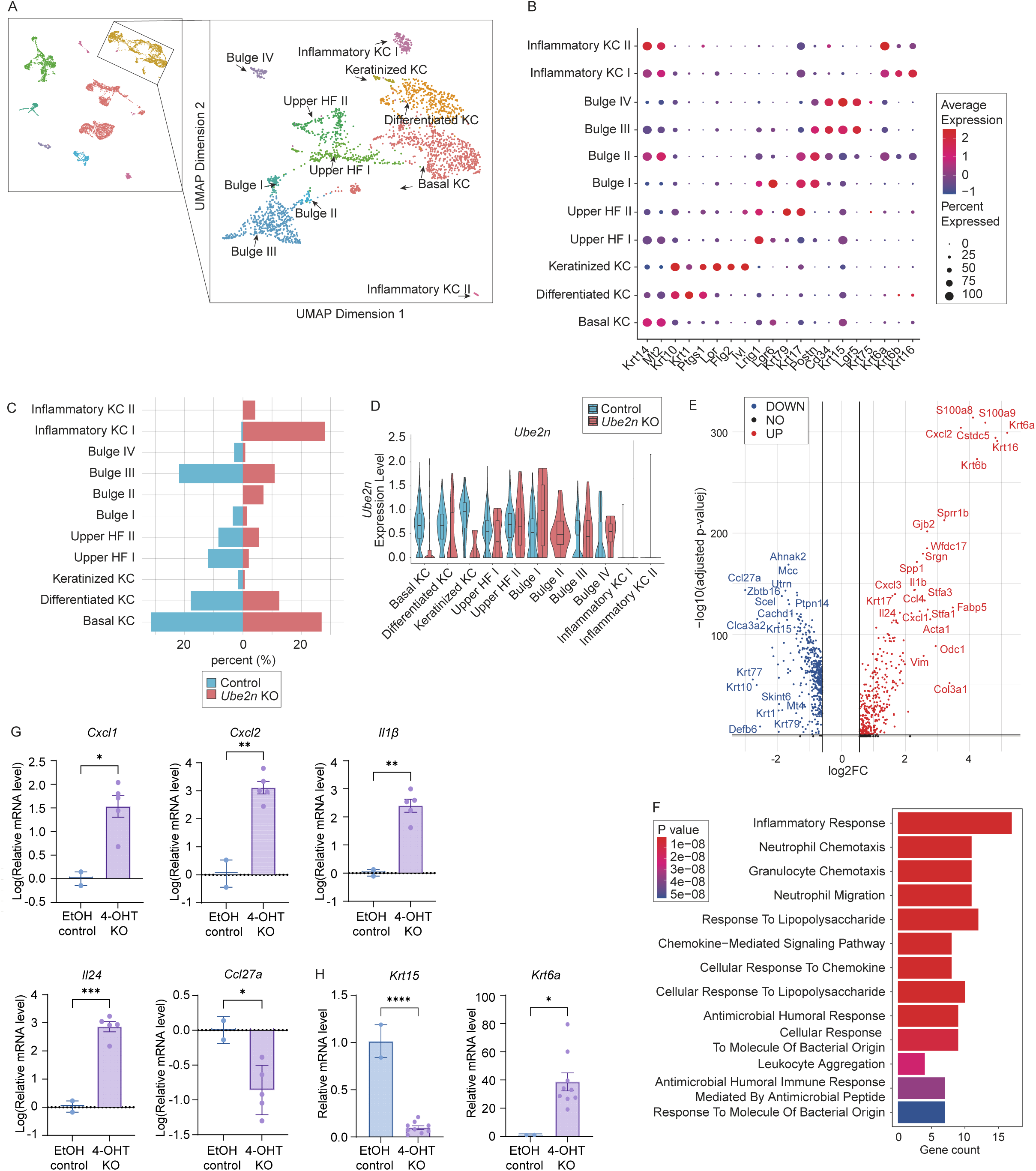
*Ube2n* KO epidermal cells display abnormal growth and differentiation and are highly inflammatory. A. UMAP of subclustering epidermal cells identifying 11 keratinocyte subgroups. B. Dot plot depicting expressions of key identifying genes. C. Percent distribution of the control and the *Ube2n* KO epidermal cells. D. Violin and bar plot of *Ube2n* expression between the control and the *Ube2n* KO group per subgroup. E. Volcano pot of differentially upregulated (red) and downregulated (blue) genes in the *Ube2n* KO epidermal cells. F. Gene ontology analysis of top 100 differentially increased genes using Biological Process 2023 reference. G. Relative mRNA expressions of *Cxcl1, Cxcl2, Il1β, Il24*, and *Ccl27a* in the mutant epidermis compared to the vehicle, ethanol (EtOH) control counterparts. Fold change is log transformed. H. Relative mRNA expressions of *Krt15* and *Krt6a* in the mutant epidermis compared to the EtOH control. *18S* is used for an internal control. * p< 0.05; ** p< 0.01; *** p < 0.001; **** p < 0.0001 (Unpaired Student’s t-test).

Keratin 6A and 6B (KRT6A/B) along with the partner Keratin 16 (KRT16) are recognized as early barrier alarmins which are often upregulated in inflammatory skin conditions such as psoriasis and actinic keratosis (37). Analysis of differentially expressed genes (DEG) between the control and the *Ube2n* KO KCs revealed 764 DEGs (adjusted p-value < 0.05) (Figure 2E). Of them, 182 DEGs were upregulated with a fold change of 2 or higher in the *Ube2n* KO KC, and 107 DEGs were downregulated (Figure 2E and Table S1). The volcano plot highlighted *Krt6a*, *Krt6b*, and *Krt16* being among the top DEGs (Figure 2E). In contrast, *Krt15* which encodes KRT15, a type I cytokeratin that is co-expressed with KRT5 throughout the basal keratinocytes including hair follicle stem cells in the bulge (38), was significantly downregulated in the KO skin, as were genes encoding suprabasal keratins such as *Krt1, Krt10, Krt77,* and *Krt79* (Figure 2E). In addition, multiple chemokines and cytokines, such as *Cxcl1, Cxcl2, Il1β*, and *Il24,* were highly upregulated, whereas *Ccl27a,* which encodes the homeostatic chemokine CCL27A, was significantly downregulated (Figure 2E). CCL27A (equivalent of a human protein CCL27) is highly expressed in normal skin but decreased in inflammatory human skin conditions including psoriasis and hidradenitis suppurativa (39–41). Gene ontology (GO) enrichment analysis of the top 100 DEGs with GO biological process reference 2023 (42) revealed activation of several prominent pathways, including inflammatory response, chemotaxis of neutrophil and other granulocytes, and neutrophil migration (Figure 2F).

In order to look more deeply into immunological changes, we utilized the gene sets of selected immunological gene ontology (GO) terms to calculate module scores based on the expression of the corresponding genes (43, 44). Specifically, we plotted the module scores for positive regulation of neutrophil chemotaxis (GO: 0090022), positive regulation of macrophage chemotaxis (GO: 0010759), and antibacterial humoral response (GO: 0019731) (42, 45, 46). We found that neutrophil chemotaxis genes were increased in the basal and differentiated KC groups as well as inflammatory KC groups of *Ube2n* KO compared to the WT control (Figures S2A-B). In contrast, macrophage chemotaxis did not show significant changes in the basal and differentiated KC groups (Figures S2C-D). The two inflammatory KC groups, however, showed increased macrophage chemotaxis genes (Figure S2C-D). Several genes among the top 100 DEGs such as *Cxcl2*, *Slpi*, and *Wfdc17* were involved in antimicrobial humoral immune responses according to the GO analysis (Figure 2F). However, the module score of antimicrobial peptides did not increase in the KO groups (Figures S2E-F), suggesting that the detected antimicrobial humoral responses may be attributed to the detection of chemokines, such as *Cxcl1* and *Cxcl2,* that regulate myeloid cell chemotaxis and migration.

To validate the scRNA-seq data, we performed RT-qPCR with RNA isolated from epidermis of *Rosa26^CreER^*.*Ube2n^fl^*^/fl^ mice. We detected significantly increased levels of *Cxcl1, Cxcl2, Il1β*, and *Il24* and a decreased level of *CCL27a* (Figure 2G). We also confirmed that the basal progenitor cell marker *Krt15* was significantly downregulated, while the inflammatory *Krt6a* was upregulated in the *Ube2n* KO epidermis (Figure 2H). These data indicate that the loss of *Ube2n* in skin cells not only leads to abnormal epidermal homeostasis but also deranged transcriptional activity of cytokines and chemokines that are involved in immune cell infiltration and progression of chronic skin inflammation.

### Deletion of UBE2N in the skin leads to increased immune infiltration and activation

To better understand the immune responses observed in the *Ube2n* KO skin, we analyzed immune cell population from the scRNA-seq data at a higher resolution. We identified 6 clusters of immune cells, including neutrophils, macrophages (MACs), dendritic cells (DCs), Langerhans cells, and lymphoid cells including T-cells and natural killer (NK) cells (Figures 3A-B). Neutrophils (*S100a8^+^*, *Cxcr2^+^*) were absent in the WT skin tissue but showed a stark increase in Ube2n KO skin (Figure 3C). There was also dramatic shift from M2-like (*Cd163^+^*, *Mrc1^+^*) to M1-like MACs (*Pdpn^+^*, *Arg1^+^*) in the KO skin (Figure 3C). Other cell types such as DCs (*Itgax^+^*, *H2-Eb1^+^*), Langerhans (*Cd207^+^*, *Epcam^+^*), and T-cells and NK cells (*Cd3g^+^*, *Nkg7^+^*) were detected in both the WT and the mutant skin (Figure 3C).

**Figure 3.**
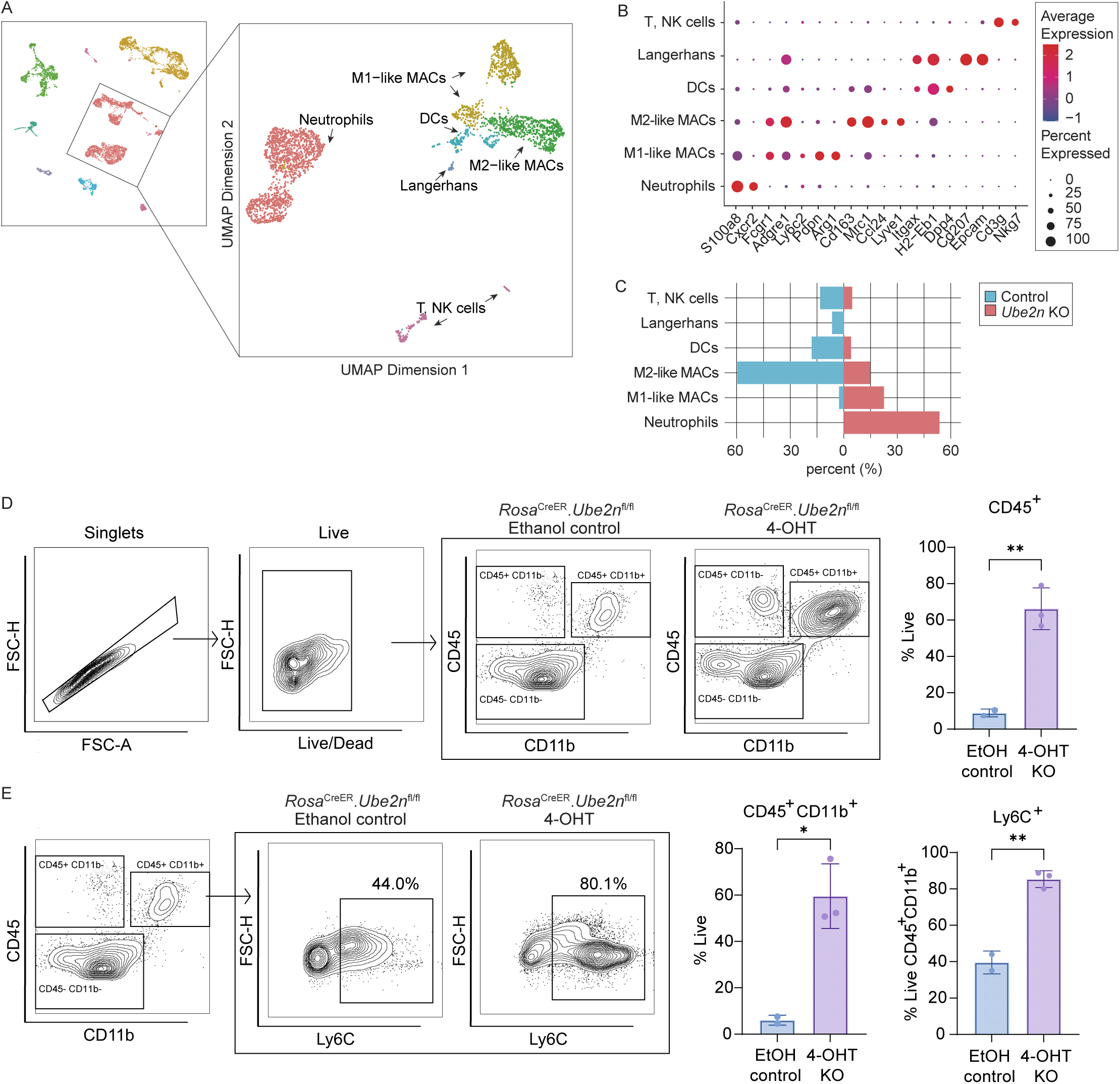
Subclustering of immune cell populations reveals myeloid cell enrichment in the *Ube2n* KO skin. A. UMAP of immune cells identify of 6 subgroups. B. Dot plot depicting expressions of key identifying genes. C. Percent distribution of different cell types of the control and the *Ube2n* KO skin. D. Gating strategy of flow cytometry analysis of *Rosa*^CreER^.*Ube2n*^fl/fl^ back skin treated with 4-OHT or EtOH vehicle control. Graph represents percent of CD45^+^ cells over live cells ± SD. E. Gating strategy of CD45^+^/CD11b^+^ myeloid cells and CD45^+^/CD11b^+^/Ly6C^+^ cells. Graphs represents percent of CD11b^+^ and Ly6C^+^ over live CD45^+^ and CD45^+^/CD11b^+^ cells, respectively ± SD. * p< 0.05; ** p< 0.01 (Unpaired Student’s t-test).

In agreement with histology and scRNA-seq data, flow cytometry detected an approximately 7-fold increase of CD45^+^ immune cells in the *Rosa26*^CreER^.*Ube2n*^fl/fl^ KO skin compared to that of the vehicle control (Figure 3D). The increased CD45^+^ cell population was mainly attributed to CD11b^+^ myeloid cells, which included CD45^+^CD11b^+^Ly6c^+^ neutrophil and monocyte populations (Figure 3E). These data indicate that loss of *Ube2n* in skin results in increased infiltration of immune cells with predominance of myeloid lineage neutrophils and macrophages.

### Deletion of UBE2N in keratinocytes is sufficient to induce skin inflammation

Data obtained with *Rosa26*^CreER^.*Ube2n*^fl/fl^ mice showed that loss of *Ube2n* is primarily localized in the epidermis. With this, we asked whether that loss of UBE2N in keratinocytes is sufficient to induce skin inflammation. To answer this question, we generated *Krt5*^CreER^.*Ube2n*^fl/fl^ mice that enabled deletion of *Ube2n* in keratin 5 (KRT5)-expressing basal keratinocytes and their subsequent progeny upon topical treatments of 4-OHT (Figure 4A). We confirmed reduced expression of UBE2N in the epidermis at the RNA level via RT-qPCR (Figure 4B) and at the protein level via western blotting (Figures 4C-D) from tissues collected 2 weeks after topical 4-OHT treatments. We found that the dorsal skin of *Krt5*^CreER^.*Ube2n*^fl/fl^ developed apparent eczematous inflammation, dry scales, and crusted erosions (Figure 4E), as well as hyperkeratosis and significantly increased epidermal and dermal thickening (Figure 4F).

**Figure 4.**
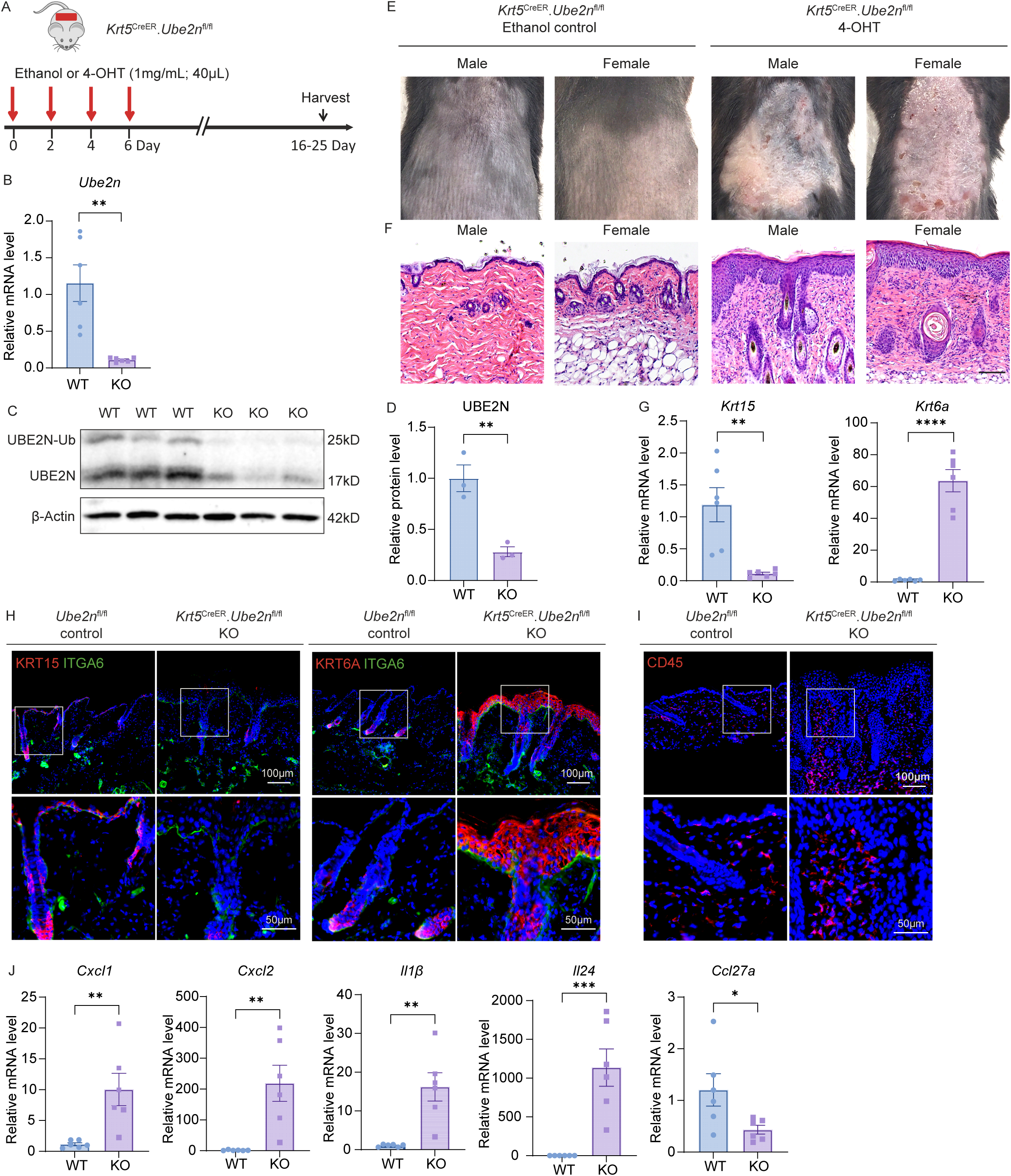
Epidermal keratinocyte-specific deletion of *Ube2n* is sufficient to induce inflammation. A. Experimental scheme of induction of *Ube2n* deletion of the *Krt5*^CreER^.*Ube2n*^flfl^ mice. B. Relative mRNA level of *Ube2n* in the *Krt5*^CreER^.*Ube2n*^flfl^ epidermis (KO) upon 4-OHT treatment compared to the *Ube2n*^fl/fl^ control (WT). C. Western blotting of protein lysates isolated from the WT and the KO epidermis. D. Quantification of western blot band intensity of UBE2N relative to β-Actin. E. Appearance of adult mice 25 days post-induction. F. H&E histological analysis of the mouse back skin sections. Scale bar: 100μm. G. Relative mRNA expressions of *Krt15* and *Krt6a* in the KO epidermis compared to the WT control mice. H. Representative immunofluorescence staining images of KRT15 (red), KRT6A (red), ITGA6 (green), and Hoechst (nuclei, blue) in the *Ube2n*^fl/fl^ control and the KO skin. Scale bars: 100μm and 50μm. I. Representative immunofluorescence staining of images of CD45 (red) and Hoechst (nuclei, blue). Scale bars: 100μm and 50μm. J. Relative mRNA expressions of *Cxcl1, Cxcl2, Il1β, Il24*, and *Ccl27a* in the KO epidermis compared to that of the *Ube2n*^fl/fl^ control. *18S* is used for internal control. * p< 0.05; ** p< 0.01; *** p < 0.001; **** p < 0.0001 (Unpaired Student’s t-test).

### Deletion of UBE2N in keratinocytes leads to changes in epidermal differentiation

To investigate UBE2N functions in epidermal homeostasis, we assessed epidermal differentiation and cell proliferation in *Krt5*^CreER^.*Ube2n*^fl/fl^ mice. We found that the basal keratinocyte progenitor cell marker KRT15 was significantly decreased in the *Ube2n* KO epidermis at both protein and mRNA levels (Figures 4G-H). In contrast, integrin α6 (ITGA6), a protein that demarcates the epidermis from the dermis, was normally expressed in the basement membrane of the mutant skin (Figures 4G-H). KRT6A was highly upregulated at both protein and mRNA level in the Ube2n KO epidermis (Figures 4G-H). Further, the UBE2N deficient epidermis was highly proliferative, as indicated by the increased number of cells expressing Ki-67, a proliferation marker shown by immunostaining and RT-qPCR (Figures S3A-B).

Next, we assessed whether immune infiltration is also increased by the loss of *Ube2n* in keratinocytes. Similar to *Rosa26*^CreER^.*Ube2n*^fl/fl^ mice, loss of *Ube2n* in *Krt5*^CreER^.*Ube2n*^fl/fl^ mice induced a significant increase of CD45^+^ immune cells in the dermis, as indicated by immunofluorescence staining (Figure 4I). Consistently, the *Ube2n* KO epidermis of the *Krt5*^CreER^.*Ube2n*^fl/fl^ mice also expressed significantly increased levels of inflammatory genes, such as *Cxcl1, Cxcl2, Il1β*, and *Il24* and a decreased level of *Ccl27a,* as measured by RT-qPCR (Figure 4J). In addition, we detected an increased level of the neutrophil marker Elastase 2 (ELA2) in the dermis (Figure S3C). These findings indicate that UBE2N expression in keratinocytes is not only essential for normal epidermal homeostasis but also crucial for maintenance of normal skin immune environment.

To further investigate the progression of the disease pathology following deletion of UBE2N, we analyzed skin changes in a time-dependent manner. Small doses of 4-OHT were applied at three different sites of the dorsal skin at different timepoints, and skin samples were collected at 7-, 11-, and 18-days (D7, D11, and D18) post first 4-OHT treatment (Figure S4A). By D18, the KO skin showed visible signs of inflammation (Figure S4B). By histological analysis, we observed that the thickening of the dermis could be seen as early as D7 and became more apparent at D11 (Figure S4C). The epidermal thickening started to show differences at D11 in some animals but became more prominent at D18 for most animals (Figure S4C). *Ube2n* mRNA expression in the epidermis was sufficiently downregulated at D11 and D18 (Figure S4D). *Krt15* mRNA expression was progressively decreased, while *Krt6a* expression was increased over the time course of the study (Figure S4E). Next, we assessed chemokine and cytokine expressions to determine their time course in the *Ube2n* KO epidermis. We detected an increase in *Cxcl1*, *Cxcl2,* and *Il1β* at the earlier timepoint (D7 and D11) in some animals, but no significant trend was observed (Figure S4F). To our surprise, the gene expression levels of the cytokines and chemokines in the epidermal cells did not show progressive changes from D11 to D18 (Figure S4F), which could be due to limited efficiency of *Ube2n* deletion. This was in contrast to the *Ube2n* KO epidermis with prolonged treatment of large doses of 4-OHT, which resulted in progressive hyperkeratosis and immune infiltration accompanied by sustained expression of cytokines and chemokines (Figure 4J). It is also possible that UBE2N regulates expression of keratins and innate immune mediators through two separate mechanisms.

### IL-1 family signaling pathways are partially responsible for inflammatory aspects of the UBE2N KO skin

The differential gene expression analysis of the scRNA-seq data revealed that *Il1β* is one of the most highly elevated genes in the *Ube2n* KO skin keratinocytes (Figure 2E). Given this, we performed more granular analysis of IL-1 family genes in different cell types. We found that *Il1β* was significantly increased in both immune cells and epithelial cells and *Il1α* was highly expressed in the immune cells of the *Ube2n* KO skin (Figure S5A). Surprisingly IL-1 receptor (*Il1r1*) expression was increased in the dermal compartments including fibroblast-like cells, endothelial cells, and smooth muscle cells in the KO skin (Figure S5A), and *Il1r2,* a competitive inhibitor of IL-1 signaling, was increased in the immune cells. Furthermore, *Il36γ*, a member of the IL-36 subgroup of IL-1 family gene, was elevated in the immune cells (Figure S5A). IL-36 signaling through IL36R in keratinocytes regulates neutrophil recruitment (47) and dysregulation of IL-36 is associated with several inflammatory skin disorders including psoriasis (48, 49). We also looked at the IL-1 signaling family genes in the epithelial and immune cell subsets. Notably, *Il1β* and *Il1r1* were significantly increased in the inflammatory KC groups, while other IL-1 signaling genes, such as *Il1α* and *Il36γ,* were highly expressed in keratinized KC (Figure S5B). *Il1β* along with *Il1α* and *Il36γ* was also highly expressed in myeloid cells including neutrophils (Figure S5C).

With these data, we hypothesized that IL-1 signaling is responsible for immunological skin lesions caused by UBE2N loss in keratinocytes. To test this hypothesis, we performed pharmacological interference of interleukin 1 receptor associated kinase 1 and 4 (IRAK1/4) which are common mediators of the TLR/IL-1 signaling pathways (50). Consistent with increased expression of IL-1 family genes, western blotting detected increased IRAK1 activation (Figure 5A), as indicated by shift of the IRAK1 band to a higher molecular weight position representing phosphorylated and ubiquitinated protein (51–53). We fed animals with chow formulated with R509, a prodrug of R835 that specifically inhibits IRAK1/4 (54), starting a week prior to the 4-OHT treatment and throughout the whole experiment (Figure 5B). Compared to mice treated with the control chow, mice treated with R509 showed decreased skin inflammation (Figure 5C). Histological analysis showed that skin of the R509 treated mice had markedly reduced hyperkeratosis and dermal thickening (Figure 5D). Phosphorylation of IRAK1 at threonine-209 (Thr 209) site initiates the activation of IRAK1 (55), and immunostaining of p-IRAK1 (Thr 209) revealed that IRAK1 activation was markedly increased in the *Ube2n* KO skin both in the epidermis as well as the dermis (Figure 5E). In addition, p-IRAK1 expressing cells were decreased in the R509 treated skin, especially in the dermis, validating the effect of the drug (Figure 5E). Immunostaining revealed reduced expression of KRT6A^+^ and decreased numbers of CD45^+^ immune cells (Figure 5F). However, KRT6A expression, albeit decreased in intensity, was still readily detectable in the IRAK1/4 inhibitor treated *Ube2n* KO skin (Figure 5F). This suggests that blocking the IL-1 signaling pathways has a potent effect on reducing immune infiltration, but it is less effective in reversing KRT6A expression in the *Ube2n* KO epidermis. Together, our data highlights a key role of UBE2N in maintenance of epidermal homeostasis and skin immunity and identify IRAK1/4 as potential therapeutic targets for inflammatory skin disorders.

**Figure 5.**
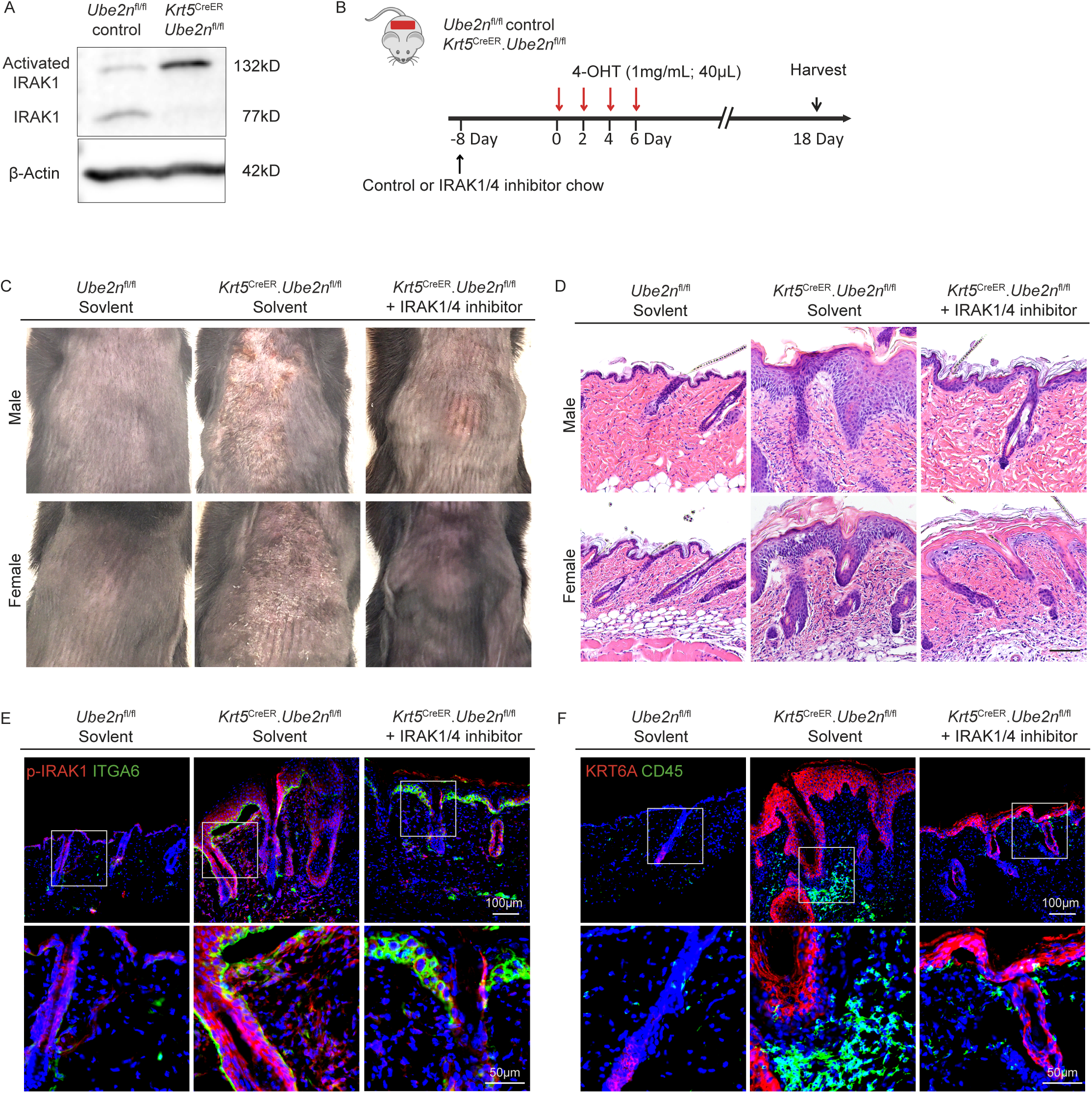
Pharmacological inhibition of IRAK1/4 alleviates skin inflammation caused by *Ube2n* deletion. A. IRAK1 protein level of the *Ube2n*^fl/fl^ control and the *Krt5*^CreER^.*Ube2n*^fl/fl^ mutant epidermis. β-Actin was used as internal control. B. Experimental scheme of induction of *Ube2n* deletion in the *Krt5*^CreER^.*Ube2n*^fl/fl^ mice and IRAK1/4 inhibition. The control and the inhibitor R509 formulated chows were provided 8 days prior to the start of 4-OHT treatment. C. Appearance of the control and the drug-treated skin analyzed at 18 days post 4-OHT treatment. D. H&E staining of the mouse back skin sections. Scale bar: 100μm. E. Representative immunofluorescence staining images of p-IRAK1 (Thr 209) (red), ITGA6 (green), and Hoechst (nuclei, blue) in the *Ube2n*^fl/fl^ control and the *Krt5*^CreER^.*Ube2n*^fl/fl^ mutant skin with or withouth IRAK1/4inhibitor. Scale bars: 100μm and 50μm. F. Representative immunofluorescence staining images of KRT6A (red), CD45 (green), and Hoechst (nuclei, blue) in the *Ube2n*^fl/fl^ control and the *Krt5*^CreER^.*Ube2n*^fl/fl^ mutant skin with or withouth IRAK1/4inhibitor. Scale bars: 100μm and 50μm.

## DISCUSSION

Our studies demonstrate that UBE2N plays a critical role in maintaining skin homeostasis. Deletion of UBE2N in adult keratinocytes leads to the development of inflammatory skin lesions characterized by hyperkeratosis, abnormal differentiation, and abundant neutrophilic immune infiltration, partially resembling skin conditions such as psoriasis and UV-damaged skin.

Our study with the *Ube2n* KO skin showed changes in keratin expression and increased expression of pro-inflammatory cytokines and myeloid-specific chemokines. The scRNA-seq analyses revealed that multiple *Ube2n* KO keratinocyte groups (basal KC, differentiated KC, inflammatory KC I, and inflammatory KC II) expressed high levels of inflammatory keratin genes such as *Krt6a*, *Krt6b*, and *Krt16*. The elevated expressions of the inflammatory keratins correlated with *Ube2n* deletion at both mRNA and protein levels and was notable as early as D7 before hyperkeratosis became apparent. This suggests that inflammatory keratin induction is an immediate downstream effect of UBE2N loss and likely occurs prior to the immune cell infiltration.

Unlike the keratin changes, chemokines and cytokine expressions showed a delayed response in the inflamed *Ube2n*-KO epidermis. There is a mild increase in *Il1β, Cxcl1*, and *Cxcl2* at the earlier time-point (D7 and D11) in some animals. However, chemokine and cytokine levels markedly changed once the skin inflammation was visibly advanced as indicated by increased hyperkeratosis and immune infiltration. Specifically, *Il1β, Cxcl1*, *Cxcl2,* and *Il24* significantly increased, while *Ccl27a* decreased. This suggests that chemokines and cytokines that recruit myeloid cells may be upregulated in the early stage of inflammation, but they may be further amplified in the later stage of inflammation by infiltrating immune cells that form a positive feedback loop with keratinocytes. Further investigation is needed to understand how these chemokines and cytokines are controlled by UBE2N.

One key skin chemokine that we found to be dysregulated by *Ube2n* KO is CCL27, a chemokine highly expressed in normal skin (36). Dysregulation of CCL27 is implicated in many inflammatory skin diseases. For example, CCL27 level is decreased in psoriasis and hidradenitis suppurativa, while its level is increased in atopic dermatitis (39–41, 56). CCL27 induces T lymphocyte recruitment, and loss of CCL27 impairs it (39). Consistent with previous findings, the *Ube2n* KO skin did not exhibit strong contribution of T lymphocytes, but rather its inflammation was primarily enriched by myeloid cells. It has been reported that the promoter region of *CCL27* harbors NF-κB binding sites and that it is regulated by p38 mitogen-activated protein kinase (MAPK) and inhibitory κB kinase (IKKβ) (57, 58). UBE2N is known to regulate both p38 and NF-κB pathways (17, 59). And therefore, it is possible that the reduced MAPK and NF-κB activities in the *Ube2n* KO epidermis may be responsible for the decreased expression of CCL27. Investigating the direct interaction between CCL27 and the MAPK or NF-κB signaling pathways could be an informative area of future research.

Among the cytokines that were dysregulated by *Ube2n* KO skin, *Il1β* was increased in both immune cells and epidermal keratinocytes as measured by scRNA-seq and RT-qPCR analyses. Through pharmacological inhibition of IRAK1/4, the key mediators of the TLR/IL-1 signaling pathways, we demonstrated that the IL-1 signaling pathway is indeed important in mediating immune infiltration and epidermal abnormalities of *Ube2n* KO skin. Blocking IL-1 activity has been used successfully to treat various autoinflammatory syndromes, such as rheumatoid arthritis, Crohn’s disease, and type 2 diabetes (60). Many of the approved therapies for blocking IL-1 activity are neutralizing antibodies or receptor antagonists (60). IRAK1/4, as common mediators of the broader TLR and IL-1 signaling pathways, have been recognized as emerging therapeutic targets for the above mentioned inflammatory diseases such as rheumatic arthritis, as well as various cancers such as myelodysplastic syndromes, leukemias, and other solid tumors (61, 62). The R835, which is the active metabolite of the pro-drug R509 used in this study, has been in a Phase 1 clinical trial for patients with lower-risk myelodysplastic syndrome (MDS) (54). Further studies may be directed to testing this agent and other IRAK1/4 inhibitors in inflammatory skin disorders such as psoriasis and radiation-damaged skin.

Interestingly, the changes in the epidermis indicated by expression of KRT6A seemed to be somewhat independent of IRAK1/4 signaling. While IRAK1/4 inhibitor treatment reduced the thickening of the epidermis, it only minimally reduced KRT6 expression in the *Ube2n* KO skin. This implies that the changes in keratin expression in keratinocytes are likely independent of IL-1 signaling pathway and that the IRAK1/4 inhibitor may act primarily on the immune cells.

Another member of the IL-1 superfamily, IL-36 signaling in keratinocyte plays a major role in neutrophil recruitment (47). IL-36 is involved in many inflammatory skin disorders including psoriasis and other systemic inflammatory diseases such as systemic lupus erythematosus (SLE), arthritis, and inflammatory bowel disease (IBD) (48, 49, 63). Upon activation by IL-36 agonists, IL-36R (also known as IL1RL2 or IL-1Rrp2) is recruited to form IL-36 receptor dimer with IL-1R which then triggers activation of the downstream NF-κB and MAPK signaling pathways (64). It has been recently reported that IL-36R undergoes K63-polyubiquitination by an E3 ligase RNF125 and the ubiquitinated IL-36R is then trafficked to lysosomes from cell surface, facilitating its turnover (64). Whether this polyubiquitination of IL-36R is mediated by UBE2N is yet to be determined. Interesting, we detected an increased level of *Il36γ* mRNA in neutrophil via scRNA-seq and in the whole skin of *Ube2n* KO animals by RT-qPCR. It has been previously shown that keratinocyte IL36R is required for neutrophilic recruitment in psoriasiform inflammation of the skin (65). It is possible that neutrophil derived IL-36γ acts on keratinocyte IL-36R to further enhance neutrophil recruitment, thereby forming a positive feedback loop.

While homozygous deletion of *Ube2n* in skin is lethal (22), dysregulation of *Ube2n* is relevant to many disease conditions. UBE2N is upregulated in cancer cells and acts in a cancer cell-intrinsic fashion to promote cell proliferation and malignancy of several cancers, such as breast cancer, neuroblastoma, colon cancer, lymphoma, and melanoma (23, 26, 27, 66–68). UBE2N plays a key role in antiviral innate immune responses mediated by the retinoic acid-inducible gene I (RIG-I)/MDA5 and mitochondrial antiviral-signaling protein (MAVS) pathway (69–71). On the other hand, UBE2N is negatively regulated by LGP2 which is a homolog and regulator of RIG-I/MDA5 (14). Of particular interest, UBE2N is itself a vulnerable target of pathogens such as *Legionella pneumophila* and *Shigella flexneri*, both of which inactivate UBE2N via covalent modification of UBE2N (72–74). While highly speculative, it is conceivable that unknown viral or bacterial pathogens deactivate UBE2N, thereby triggering the onset of inflammatory skin reactions such as psoriasis. Lastly, whether our mouse models of the conditional *Ube2n* KO in the skin lead to dysregulated antimicrobial defense or changes in skin microbiota is yet to be determined and could be a very interesting area of research.

In summary, our findings underscore that UBE2N suppresses inflammation in the skin, and its loss in keratinocyte populations leads to progressive skin inflammation with increased expression of pro-inflammatory cytokines and chemokines that favor myeloid infiltration. UBE2N also maintains the homeostasis of epidermal proliferation and differentiation, and its loss leads to hyperkeratosis accompanied by disrupted epidermal differentiation and loss of epidermal stem cell marker expression. Our findings suggest that IRAK1/4 can be a potential therapeutic target for inflammatory skin disorders associated with abnormal myeloid infiltration, such as pustular psoriasis and neutrophilic dermatosis. Together, our data establish a novel inflammatory axis in the skin, showing that UBE2N control of inflammatory skin disorders may be rectified through IRAK1/4 modulation.

## MATERIALS AND METHODS

### Mouse models

Animal studies were conducted in accordance with protocols approved by Duke Animal Care and Use Committee. All mice were maintained under a temperature controlled and specific pathogen-free environment. *Krt5*^CreERT2^ (strain #: 029155; (75)) was obtained from Jackson laboratories (Bar Harbor, ME). *Rosa26*^CreER^.*Ube2n*^fl/fl^ mice were kindly gifted by Dr. Shao-Cong Sun (University of Texas MD Anderson, Houston, TX) under permission of Dr. Shizuo Akira (Osaka University, Osaka, Japan) (17). The *Krt5*^CreER^.*Ube2n*^fl/fl^ was generated via crossbreeding and genotyped via PCR with primers listed in Table S2. Both male and female mice were used in this study. To induce local deletion of *Ube2n* in the skin, 1-6 months-old adult mice were shaved on the dorsal skin and treated topically with 40 μl of 1-5mg/ml 4-hydroxytamoxifen (4-OHT; Sigma, St. Louis, MO) in ethanol every other day for 4 times. For the time-course experiment, 20 μl of 1 mg/ml of 4-OHT was applied daily for 4 days. The vehicle control mice were treated with ethanol. For the WT control, littermates *Ube2n*^fl/fl^ mice were treated with equal amounts of 4-OHT.

### Skin tissue collection

The back skin tissue was collected and placed in OCT (Sakura Finetek, Torrence, CA) for histology or immunofluorescence analysis. For epidermal and dermal separation, the skin was incubated with dermal side down on 2.5mg/mL of dispase II (ThermoFisher, Waltham, MA) in PBS at 4°C overnight. The epidermis was gently separated with forceps. The tissues were chopped and harvested in TRI Reagent (Zymo Research, Tustin, CA) or Trizol reagent (ThermoFisher) and kept at −80°C for RNA extraction or in RIPA buffer containing protease inhibitor, phosphatase inhibitor, 5mM EDTA (ThermoFisher), 5uM N-Ethylmaleimide (NEM; Sigma), 1mM phenylmethylsulfonyl fluoride (PMSF, Sigma), 1mM dithiothreitol (DTT, Sigma), 5mM iodoacetamide (IAA, UBPBio, Dallas, TX), 1mM sodium orthovanadate (Na3VO4, Sigma), and 1uM ubiquitin-aldehyde (South Bay Bio, San Jose, CA).

### Mouse skin single cell isolation

1×8mm punch biopsy or 2×4mm punch biopsies were used to isolate single cells. The tissues were washed with RPMI (ThermoFisher) on ice. The tissues were cut into small pieces with scissors and incubated in 3mL of RPMI containing 900µg of Liberase TM (Roche, Basal, Switzerland) and 150µg DNase I (MP Biomedicals, Santa Ana, CA) at 37°C for 30 minutes and gently shaken every 10 minutes. Cells were spun down at 350g for 6 minutes and were re-suspended with 4mL of 0.05% Trypsin (ThermoFisher) and incubated at 37°C for 10 minutes. Cells were filtered through 40µm cell strainer with 4mL of RPMI + 5% FBS (R&D Systems, Minneapolis, MN), and the unfiltered tissues were further digested with 4mL of 0.05% Trypsin and incubated at 37°C for 10 minutes. Cells were filtered again with 4mL of RPMI + 5% FBS. The cell suspension was spun down at 350g for 6 minutes. The pellets were washed twice with 5ml appropriate buffers: for scRNA-seq, PBS containing 0.04% BSA (VWR, Radnor, PA) and for FACS, PBS containing 2% FBS and 2mM EDTA in PBS.

### Single-cell RNA-seq extraction and library preparation

After quality check, the cell suspensions prepared above were immediately sent to Duke Molecular Genomics core for barcoding, cDNA library construction, and subsequent sequencing on the NovaSeq6000 Illumina sequencing platform at 50K reads/cell. The data is deposited in NCBI (tracking #2439013) with GEO accession number to be generated.

### Single-cell RNA-sequence analysis workflow

The primary analytical pipeline for the scRNA-seq analysis followed the recommended protocols from 10X Genomics. Briefly, we demultiplexed the generated raw base call (BCL) files into FASTQ files. CellRanger (version 7.1.0) count function was used to align the reads to the mm10 mouse reference genome and generate the cell-gene count matrix following filtering unviable cells based on low number of unique RNA molecules per cell. The filtered count matrix was read using Seurat (version 4.1.1) (76) on R (version 4.0.5). Seurat was used to create a Seurat object from the read count matrix. Subsequent quality control, exploratory, and differential expression analyses were also done using Seurat. Cells were further filtered based on the number of unique RNA molecules observed (minimum of 250 and maximum of 45000), percent of counts ascribed to mitochondrial genes (maximum of 12%), the number of detected genes per cell (minimum of 600 genes and maximum of 7500). The thresholds were used to prune out outliers based on an initial plot of these quality control metrics before filtering—to exclude low quality cells and possible doublets. The counts were then normalized for sequencing depth using the SCTransform method (77) during which the percentage of mitochondrial counts was used as a regression variable. Cells from the various batches of sequencing were then integrated using the Seurat anchor method (78) using the top 3000 variable features and with SCTransform as the normalization method. Subsequently, clustering (resolution of 1.0) and UMAP were done using 100 principal components. Identification of cell types as well as the choice of optimal clustering parameters were guided by the localization of cell type-specific genes. Marker genes used for each cell type have been shown in the corresponding figures in this manuscript. Sub-clustering of epithelial cells and immune cells (resolution of 0.8) and UMAP were done using 30 principal components. For differential expression analysis, a false discovery rate (also known as adjusted p-value) of less than 0.05 and a log2foldchange of 0.58 were used as thresholds to call significantly dysregulated genes. EnrichR (79–81) (version 3.2) was used for gene ontology enrichment analysis and for plotting the enrichment plots. GO terms and associated gene lists used for calculating module score were obtained from Gene Ontology Browser tool from the Mouse Genome Informatics (https://www.informatics.jax.org/vocab/gene_ontology/).

### Flow cytometry

Antibodies used in the study include CD45 (30-F11), Ly6C (HK1.4), CD11b (M1/70). Antibodies were conjugated to FITC, PerCP-Cy5.5, PE-Cy7. Antibody information can be found in Table S2. Ghost Dye Violet 510 was used for viability (Tonbo Biosciences, San Diego, CA). Cells were captured with BD FACSDiva software (BD Biosciences, Franklin Lakes, NJ) on BD FACSCanto (BD Biosciences) analyzed with FlowJo software (FlowJo LLC, Ashland, OR).

### RNA extraction and RT-qPCR

RNA was extracted using the Direct-zol RNA Purification Kit (Zymo Research). RNA was transcribed into cDNA using the iScript cDNA synthesis kit (BioRad, Hercules, CA). The cDNA was used for RT-qPCR with qPCRBIO SyGreen Blue mix Hi-ROX (PCR Biosystems, London, UK) on the StepOne Plus Real-Time PCR machine (Applied Biosystems, Foster City, CA) and primers listed in Table S2. Fold change of gene expression was calculated and normalized to the housekeeping gene *18S* and calculated using the 2(-ΔΔCt) method (82).

### H&E histology

Hematoxylin and eosin staining was performed in Duke Pathology lab (Durham, NC). Frozen tissue in OCT was thawed and paraffin embedded. The skin tissues were sliced into 9µm sections. Sections were stained with hematoxylin (Cancer Diagnostics, Durham NC) and eosin (Cancer Diagnostics). The sections were imaged with an Olympus BX41 microscope (Olympus, Tokyo, Japan).

### Immunofluorescent staining

Mouse skin was frozen in OCT for cryosectioning. 9µm sections were fixed in 4% paraformaldehyde, washed in PBS, and permeabilized in 0.1% Triton X-100 (10 minutes) and blocked in a blocking buffer containing 2.5% normal goat serum, 2.5% normal donkey serum 1% BSA, 2% gelatin and 0.01% Triton X-100 (1 hour). The sections were then incubated overnight with primary antibodies (4 °C): KRT15, KRT6A, ITGA6, CD45, Ki-67, ELA2, p-IRAK1 (Thr 209), rabbit IgG, and rat IgG as a control. Antibody information can be found in Table S2. Followed by washing, appropriate secondary antibodies were treated for 1-2 hours in the dark (AF555 anti-Rabbit, AF555 anti-Rat, AF647 anti-Rat (Invitrogen, Waltham, MA)) and washed in PBS containing 0.01% Triton X-100. Sections were counterstained with Hoechst 33342 (Invitrogen) and mounted using ProLong Gold antifade reagent (Invitrogen). Cells were imaged using the Olympus Cell Sens software (Olympus).

### Western Blot

Whole cell lysates were quantified using DC protein assay (BioRad). Equal amounts of protein were separated on SDS–PAGE 12% gels, transferred onto PVDF membranes, and blocked with 5% BSA (VWR). Membranes were incubated with UBE2N, IRAK, or β-Actin overnight at 4°C. Antibody information can be found in Table S2. The antibodies were detected with anti-rabbit IgG IRDye 680RD (LI-COR, Lincoln, NE) and imaged using Odyssey Fc Imaging System (LI-COR). Quantification analysis was performed using Image Lab software (BioRad).

### Statistical analysis and graphs

Figures and statistical analyses were conducted using Graphpad Prism software (GraphPad, San Diego, CA). Figures were created with means ± standard error of the mean (SEM). Statistical analyses were performed using the two-tailed unpaired Student’s t test or two-way ANOVA.

## ACKNOWLEDGEMENTS

This study was supported by funding from NIH/NIAMS 5R01AR073858 to JZ, and the Department of Dermatology of Duke University. We thank Dr. Shizuo Akira of Osaka University and Dr. Shao-Cong Sun of University of Texas MD Anderson for providing the *Rosa*^CreER^.*Ube2n*^fl/fl^ mice. We also thank Rigel Pharmaceuticals for providing the IRAK1/4 inhibitor. We thank Raphael Lee for technical assistance.

## AUTHOR CONTRIBUTIONS

ML, MH, and JZ conceptualized the study. ML, MH, WM, AO, YJ, HS, VJ, VM, and JZ contributed to methodology. ML, MH, WM, YJ, HS, and JZ performed animal studies and analyzed the data. ML, AO, VJ, YD, SG, and JZ performed bioinformatic analysis. ML and JZ wrote the manuscript and prepared the figures. All authors reviewed and edited the manuscript.

## REFERENCES

1 Liao Y, Sumara I, Pangou E. Non-proteolytic ubiquitylation in cellular signaling and human disease. Commun Biol. 2022;5(1):114.

2 Akutsu M, Dikic I, Bremm A. Ubiquitin chain diversity at a glance. J Cell Sci. 2016;129(5):875–80.

3 French ME, Koehler CF, Hunter T. Emerging functions of branched ubiquitin chains. Cell Discov. 2021;7(1):6.

4 Sun SC. CYLD: a tumor suppressor deubiquitinase regulating NF-kappaB activation and diverse biological processes. Cell Death Differ. 2010;17(1):25–34.

5 Komander D, Lord CJ, Scheel H, Swift S, Hofmann K, Ashworth A, et al. The structure of the CYLD USP domain explains its specificity for Lys63-linked polyubiquitin and reveals a B box module. Mol Cell. 2008;29(4):451–64.

6 Soss SE, Yue Y, Dhe-Paganon S, Chazin WJ. E2 conjugating enzyme selectivity and requirements for function of the E3 ubiquitin ligase CHIP. J Biol Chem. 2011;286(24):21277–86.

7 Mattiroli F, Vissers JH, van Dijk WJ, Ikpa P, Citterio E, Vermeulen W, et al. RNF168 ubiquitinates K13-15 on H2A/H2AX to drive DNA damage signaling. Cell. 2012;150(6):1182–95.

8 Hodge CD, Spyracopoulos L, Glover JN. Ubc13: the Lys63 ubiquitin chain building machine. Oncotarget. 2016;7(39):64471–504.

9 Deng L, Wang C, Spencer E, Yang L, Braun A, You J, et al. Activation of the IkappaB kinase complex by TRAF6 requires a dimeric ubiquitin-conjugating enzyme complex and a unique polyubiquitin chain. Cell. 2000;103(2):351–61.

10 Chung JY, Park YC, Ye H, Wu H. All TRAFs are not created equal: common and distinct molecular mechanisms of TRAF-mediated signal transduction. J Cell Sci. 2002;115(Pt 4):679–88.

11 Kobayashi T, Walsh MC, Choi Y. The role of TRAF6 in signal transduction and the immune response. Microbes Infect. 2004;6(14):1333–8.

12 Xia ZP, Sun L, Chen X, Pineda G, Jiang X, Adhikari A, et al. Direct activation of protein kinases by unanchored polyubiquitin chains. Nature. 2009;461(7260):114-9.

13 Lamothe B, Besse A, Campos AD, Webster WK, Wu H, Darnay BG. Site-specific Lys-63-linked tumor necrosis factor receptor-associated factor 6 auto-ubiquitination is a critical determinant of I kappa B kinase activation. J Biol Chem. 2007;282(6):4102–12.

14 Lenoir JJ, Parisien JP, Horvath CM. Immune regulator LGP2 targets Ubc13/UBE2N to mediate widespread interference with K63 polyubiquitination and NF-kappaB activation. Cell Rep. 2021;37(13):110175.

15 Hofmann RM, Pickart CM. In vitro assembly and recognition of Lys-63 polyubiquitin chains. J Biol Chem. 2001;276(30):27936–43.

16 Hofmann RM, Pickart CM. Noncanonical MMS2-encoded ubiquitin-conjugating enzyme functions in assembly of novel polyubiquitin chains for DNA repair. Cell. 1999;96(5):645–53.

17 Yamamoto M, Okamoto T, Takeda K, Sato S, Sanjo H, Uematsu S, et al. Key function for the Ubc13 E2 ubiquitin-conjugating enzyme in immune receptor signaling. Nat Immunol. 2006;7(9):962–70.

18 Beck DB, Ferrada MA, Sikora KA, Ombrello AK, Collins JC, Pei W, et al. Somatic Mutations in UBA1 and Severe Adult-Onset Autoinflammatory Disease. N Engl J Med. 2020;383(27):2628–38.

19 Erpapazoglou Z, Walker O, Haguenauer-Tsapis R. Versatile roles of k63-linked ubiquitin chains in trafficking. Cells. 2014;3(4):1027–88.

20 Wooff J, Pastushok L, Hanna M, Fu Y, Xiao W. The TRAF6 RING finger domain mediates physical interaction with Ubc13. FEBS Lett. 2004;566(1-3):229–33.

21 Ye Y, Rape M. Building ubiquitin chains: E2 enzymes at work. Nat Rev Mol Cell Biol. 2009;10(11):755–64.

22 Sayama K, Yamamoto M, Shirakata Y, Hanakawa Y, Hirakawa S, Dai X, et al. E2 Polyubiquitin-conjugating enzyme Ubc13 in keratinocytes is essential for epidermal integrity. J Biol Chem. 2010;285(39):30042–9.

23 Cheng J, Fan YH, Xu X, Zhang H, Dou J, Tang Y, et al. A small-molecule inhibitor of UBE2N induces neuroblastoma cell death via activation of p53 and JNK pathways. Cell Death Dis. 2014;5(2):e1079.

24 Barreyro L, Sampson AM, Ishikawa C, Hueneman KM, Choi K, Pujato MA, et al. Blocking UBE2N abrogates oncogenic immune signaling in acute myeloid leukemia. Sci Transl Med. 2022;14(635):eabb7695.

25 Wu Z, Shen S, Zhang Z, Zhang W, Xiao W. Ubiquitin-conjugating enzyme complex Uev1A-Ubc13 promotes breast cancer metastasis through nuclear factor-small ka, CyrillicB mediated matrix metalloproteinase-1 gene regulation. Breast Cancer Res. 2014;16(4):R75.

26 Gombodorj N, Yokobori T, Yoshiyama S, Kawabata-Iwakawa R, Rokudai S, Horikoshi I, et al. Inhibition of Ubiquitin-conjugating Enzyme E2 May Activate the Degradation of Hypoxia-inducible Factors and, thus, Overcome Cellular Resistance to Radiation in Colorectal Cancer. Anticancer Res. 2017;37(5):2425–36.

27 Dikshit A, Jin YJ, Degan S, Hwang J, Foster MW, Li CY, et al. UBE2N Promotes Melanoma Growth via MEK/FRA1/SOX10 Signaling. Cancer Res. 2018;78(22):6462–72.

28 Dikshit A, Zhang JY. UBE2N plays a pivotal role in maintaining melanoma malignancy. Oncotarget. 2018;9(100):37347–8.

29 Alameda JP, Fernandez-Acenero MJ, Moreno-Maldonado R, Navarro M, Quintana R, Page A, et al. CYLD regulates keratinocyte differentiation and skin cancer progression in humans. Cell Death Dis. 2011;2:e208.

30 Oudot T, Lesueur F, Guedj M, de Cid R, McGinn S, Heath S, et al. An association study of 22 candidate genes in psoriasis families reveals shared genetic factors with other autoimmune and skin disorders. J Invest Dermatol. 2009;129(11):2637–45.

31 Miliani de Marval P, Lutfeali S, Jin JY, Leshin B, Selim MA, Zhang JY. CYLD inhibits tumorigenesis and metastasis by blocking JNK/AP1 signaling at multiple levels. Cancer Prev Res (Phila). 2011;4(6):851–9.

32 Jin YJ, Wang S, Cho J, Selim MA, Wright T, Mosialos G, et al. Epidermal CYLD inactivation sensitizes mice to the development of sebaceous and basaloid skin tumors. JCI Insight. 2016;1(11).

33 Agrawal R, Woodfolk JA. Skin barrier defects in atopic dermatitis. Curr Allergy Asthma Rep. 2014;14(5):433.

34 Orsmond A, Bereza-Malcolm L, Lynch T, March L, Xue M. Skin Barrier Dysregulation in Psoriasis. Int J Mol Sci. 2021;22(19).

35 Garelli CJ, Refat MA, Nanaware PP, Ramirez-Ortiz ZG, Rashighi M, Richmond JM. Current Insights in Cutaneous Lupus Erythematosus Immunopathogenesis. Front Immunol. 2020;11:1353.

36 Davila ML, Xu M, Huang C, Gaddes ER, Winter L, Cantorna MT, et al. CCL27 is a crucial regulator of immune homeostasis of the skin and mucosal tissues. iScience. 2022;25(6):104426.

37 Zhang X, Yin M, Zhang LJ. Keratin 6, 16 and 17-Critical Barrier Alarmin Molecules in Skin Wounds and Psoriasis. Cells. 2019;8(8).

38 Bose A, Teh MT, Mackenzie IC, Waseem A. Keratin k15 as a biomarker of epidermal stem cells. Int J Mol Sci. 2013;14(10):19385–98.

39 Homey B, Alenius H, Muller A, Soto H, Bowman EP, Yuan W, et al. CCL27-CCR10 interactions regulate T cell-mediated skin inflammation. Nat Med. 2002;8(2):157–65.

40 Gudjonsson JE, Ding J, Johnston A, Tejasvi T, Guzman AM, Nair RP, et al. Assessment of the psoriatic transcriptome in a large sample: additional regulated genes and comparisons with in vitro models. J Invest Dermatol. 2010;130(7):1829–40.

41 Hotz C, Boniotto M, Guguin A, Surenaud M, Jean-Louis F, Tisserand P, et al. Intrinsic Defect in Keratinocyte Function Leads to Inflammation in Hidradenitis Suppurativa. J Invest Dermatol. 2016;136(9):1768–80.

42 Thomas PD, Ebert D, Muruganujan A, Mushayahama T, Albou LP, Mi H. PANTHER: Making genome-scale phylogenetics accessible to all. Protein Sci. 2022;31(1):8–22.

43 Hao Y, Hao S, Andersen-Nissen E, Mauck WM, 3rd, Zheng S, Butler A, et al. Integrated analysis of multimodal single-cell data. Cell. 2021;184(13):3573–87 e29.

44 Tirosh I, Izar B, Prakadan SM, Wadsworth MH, 2nd, Treacy D, Trombetta JJ, et al. Dissecting the multicellular ecosystem of metastatic melanoma by single-cell RNA-seq. Science. 2016;352(6282):189–96.

45 Ashburner M, Ball CA, Blake JA, Botstein D, Butler H, Cherry JM, et al. Gene ontology: tool for the unification of biology. The Gene Ontology Consortium. Nat Genet. 2000;25(1):25–9.

46 Gene Ontology C, Aleksander SA, Balhoff J, Carbon S, Cherry JM, Drabkin HJ, et al. The Gene Ontology knowledgebase in 2023. Genetics. 2023;224(1).

47 Koss CK, Wohnhaas CT, Baker JR, Tilp C, Przibilla M, Lerner C, et al. IL36 is a critical upstream amplifier of neutrophilic lung inflammation in mice. Commun Biol. 2021;4(1):172.

48 Tortola L, Rosenwald E, Abel B, Blumberg H, Schafer M, Coyle AJ, et al. Psoriasiform dermatitis is driven by IL-36-mediated DC-keratinocyte crosstalk. J Clin Invest. 2012;122(11):3965–76.

49 Buhl AL, Wenzel J. Interleukin-36 in Infectious and Inflammatory Skin Diseases. Front Immunol. 2019;10:1162.

50 Jain A, Kaczanowska S, Davila E. IL-1 Receptor-Associated Kinase Signaling and Its Role in Inflammation, Cancer Progression, and Therapy Resistance. Front Immunol. 2014;5:553.

51 Yamin TT, Miller DK. The interleukin-1 receptor-associated kinase is degraded by proteasomes following its phosphorylation. J Biol Chem. 1997;272(34):21540–7.

52 Pauls E, Nanda SK, Smith H, Toth R, Arthur JSC, Cohen P. Two phases of inflammatory mediator production defined by the study of IRAK2 and IRAK1 knock-in mice. J Immunol. 2013;191(5):2717–30.

53 Vollmer S, Strickson S, Zhang T, Gray N, Lee KL, Rao VR, et al. The mechanism of activation of IRAK1 and IRAK4 by interleukin-1 and Toll-like receptor agonists. Biochem J. 2017;474(12):2027–38.

54 Lamagna C, Gundel C, Chan M, Young C, Braselmann S, Frances R, et al. FRI0016 R835, A NOVEL IRAK1/4 DUAL INHIBITOR IN CLINICAL DEVELOPMENT, BLOCKS TOLL-LIKE RECEPTOR 4 (TLR4) SIGNALING IN HUMAN AND MOUSE. Annals of the Rheumatic Diseases. 2020;79(Suppl 1):579-.

55 Kollewe C, Mackensen AC, Neumann D, Knop J, Cao P, Li S, et al. Sequential autophosphorylation steps in the interleukin-1 receptor-associated kinase-1 regulate its availability as an adapter in interleukin-1 signaling. J Biol Chem. 2004;279(7):5227–36.

56 Kakinuma T, Saeki H, Tsunemi Y, Fujita H, Asano N, Mitsui H, et al. Increased serum cutaneous T cell-attracting chemokine (CCL27) levels in patients with atopic dermatitis and psoriasis vulgaris. J Allergy Clin Immunol. 2003;111(3):592–7.

57 Vestergaard C, Johansen C, Otkjaer K, Deleuran M, Iversen L. Tumor necrosis factor-alpha-induced CTACK/CCL27 (cutaneous T-cell-attracting chemokine) production in keratinocytes is controlled by nuclear factor kappaB. Cytokine. 2005;29(2):49–55.

58 Riis JL, Johansen C, Vestergaard C, Otkjaer K, Kragballe K, Iversen L. CCL27 expression is regulated by both p38 MAPK and IKKbeta signalling pathways. Cytokine. 2011;56(3):699–707.

59 Chang JH, Xiao Y, Hu H, Jin J, Yu J, Zhou X, et al. Ubc13 maintains the suppressive function of regulatory T cells and prevents their conversion into effector-like T cells. Nat Immunol. 2012;13(5):481–90.

60 Dinarello CA, Simon A, van der Meer JW. Treating inflammation by blocking interleukin-1 in a broad spectrum of diseases. Nat Rev Drug Discov. 2012;11(8):633–52.

61 Bennett J, Starczynowski DT. IRAK1 and IRAK4 as emerging therapeutic targets in hematologic malignancies. Curr Opin Hematol. 2022;29(1):8–19.

62 Singer JW, Fleischman A, Al-Fayoumi S, Mascarenhas JO, Yu Q, Agarwal A. Inhibition of interleukin-1 receptor-associated kinase 1 (IRAK1) as a therapeutic strategy. Oncotarget. 2018;9(70):33416–39.

63 Yuan ZC, Xu WD, Liu XY, Liu XY, Huang AF, Su LC. Biology of IL-36 Signaling and Its Role in Systemic Inflammatory Diseases. Front Immunol. 2019;10:2532.

64 Saha SS, Caviness G, Yi G, Raymond EL, Mbow ML, Kao CC. E3 Ubiquitin Ligase RNF125 Activates Interleukin-36 Receptor Signaling and Contributes to Its Turnover. J Innate Immun. 2018;10(1):56–69.

65 Hernandez-Santana YE, Leon G, St Leger D, Fallon PG, Walsh PT. Keratinocyte interleukin-36 receptor expression orchestrates psoriasiform inflammation in mice. Life Sci Alliance. 2020;3(4).

66 Wu X, Zhang W, Font-Burgada J, Palmer T, Hamil AS, Biswas SK, et al. Ubiquitin-conjugating enzyme Ubc13 controls breast cancer metastasis through a TAK1-p38 MAP kinase cascade. Proc Natl Acad Sci U S A. 2014;111(38):13870–5.

67 Pulvino M, Liang Y, Oleksyn D, DeRan M, Van Pelt E, Shapiro J, et al. Inhibition of proliferation and survival of diffuse large B-cell lymphoma cells by a small-molecule inhibitor of the ubiquitin-conjugating enzyme Ubc13-Uev1A. Blood. 2012;120(8):1668–77.

68 Wu Z, Shen S, Zhang Z, Zhang W, Xiao W. Ubiquitin-conjugating enzyme complex Uev1A-Ubc13 promotes breast cancer metastasis through nuclear factor-кB mediated matrix metalloproteinase-1 gene regulation. Breast cancer research : BCR. 2014;16(4):R75.

69 Fletcher AJ, Mallery DL, Watkinson RE, Dickson CF, James LC. Sequential ubiquitination and deubiquitination enzymes synchronize the dual sensor and effector functions of TRIM21. Proc Natl Acad Sci U S A. 2015;112(32):10014–9.

70 Shi Y, Yuan B, Zhu W, Zhang R, Li L, Hao X, et al. Ube2D3 and Ube2N are essential for RIG-I-mediated MAVS aggregation in antiviral innate immunity. Nat Commun. 2017;8:15138.

71 Kiss L, Zeng J, Dickson CF, Mallery DL, Yang JC, McLaughlin SH, et al. A tri-ionic anchor mechanism drives Ube2N-specific recruitment and K63-chain ubiquitination in TRIM ligases. Nat Commun. 2019;10(1):4502.

72 Puvar K, Iyer S, Fu J, Kenny S, Negron Teron KI, Luo ZQ, et al. Legionella effector MavC targets the Ube2N∼Ub conjugate for noncanonical ubiquitination. Nat Commun. 2020;11(1):2365.

73 Gan N, Nakayasu ES, Hollenbeck PJ, Luo ZQ. Legionella pneumophila inhibits immune signalling via MavC-mediated transglutaminase-induced ubiquitination of UBE2N. Nat Microbiol. 2019;4(1):134–43.

74 Nishide A, Kim M, Takagi K, Himeno A, Sanada T, Sasakawa C, et al. Structural basis for the recognition of Ubc13 by the Shigella flexneri effector OspI. J Mol Biol. 2013;425(15):2623–31.

75 Van Keymeulen A, Rocha AS, Ousset M, Beck B, Bouvencourt G, Rock J, et al. Distinct stem cells contribute to mammary gland development and maintenance. Nature. 2011;479(7372):189-93.

76 Satija R, Farrell JA, Gennert D, Schier AF, Regev A. Spatial reconstruction of single-cell gene expression data. Nat Biotechnol. 2015;33(5):495–502.

77 Hafemeister C, Satija R. Normalization and variance stabilization of single-cell RNA-seq data using regularized negative binomial regression. Genome Biol. 2019;20(1):296.

78 Stuart T, Butler A, Hoffman P, Hafemeister C, Papalexi E, Mauck WM, 3rd, et al. Comprehensive Integration of Single-Cell Data. Cell. 2019;177(7):1888–902 e21.

79 Chen EY, Tan CM, Kou Y, Duan Q, Wang Z, Meirelles GV, et al. Enrichr: interactive and collaborative HTML5 gene list enrichment analysis tool. BMC Bioinformatics. 2013;14:128.

80 Kuleshov MV, Jones MR, Rouillard AD, Fernandez NF, Duan Q, Wang Z, et al. Enrichr: a comprehensive gene set enrichment analysis web server 2016 update. Nucleic Acids Res. 2016;44(W1):W90–7.

81 Xie Z, Bailey A, Kuleshov MV, Clarke DJB, Evangelista JE, Jenkins SL, et al. Gene Set Knowledge Discovery with Enrichr. Curr Protoc. 2021;1(3):e90.

82 Livak KJ, Schmittgen TD. Analysis of Relative Gene Expression Data Using Real-Time Quantitative PCR and the 2−ΔΔCT Method. Methods. 2001;25(4):402–8.

